# RNA-Binding Proteins MSI-1 (Musashi) and EXC-7 (HuR) Regulate Serotonin-Mediated Behaviors in *C. elegans*

**DOI:** 10.1101/748509

**Authors:** Zhe Yang, Lan Lan, Xiaoqing Wu, Liang Xu, Matthew Buechner

## Abstract

The evolutionarily conserved RNA-binding proteins HuR and MSI are essential for multiple developmental processes and are upregulated in many cancer tissues. The *C. elegans* homologues EXC-7 (HuR) and MSI-1 (MSI1 and MSI2) have been implicated in tubulogenesis, neural development, and specific behaviors that include male tail-curling to maintain contact with the hermaphrodite during mating. This behavior is mediated by serotonin signaling. Here, drug studies plus biochemical and genetic results indicate that MSI-1 affects serotonergic signaling through stabilization of mRNA of the Gα protein GOA-1/GNAO1 in neurons, which in turn affects activity of the serotonin synthase TPH-1/tryptophan hydroxylase via the response element CRH-1/CREB. EXC-7 (HuR) is also involved in this regulatory pathway. These results indicate a novel pathway and role for these RNA-binding proteins in regulating neurotransmitter levels that could be conserved in other tissues where these RNA-binding proteins are present.

**Impact Statement**

RNA-binding proteins Musashi and HuR upregulate serotonin levels for male-specific movement during mating via a novel pathway involving a neural Gα protein, response element, and serotonin synthase.

## Introduction

RNA-binding proteins (RBP) play vital roles in the translational regulation of gene expression, mRNA distribution, stability, and degradation [1, 2]. The Hu/ELAV and Musashi families of RNA-binding proteins have been particularly well studied. The ELAV family (from *Drosophila* Embryonic Lethal Abnormal Vision [3]) has four members in humans: HuB, HuC, HuD, and HuR [4]; and one nematode homologue, EXC-7 [5, 6]. These proteins are critical for neural development, and affect stability and splicing of thousands of different mRNAs by binding to 3’ UTRs (untranslated regions) of regulated mRNAs. In mammals, HuR is highly expressed in multiple human cancers [7–9]. Several small-molecule compounds disrupt HuR-mRNA interactions [10] and slow growth of cancerous tissues. HuR increases stability of many mRNAs, including mRNAs encoding the Musashi RNA-binding proteins [11]. Similarly, in *C. elegans*, EXC-7 regulates stability of multiple mRNAs, for example that of *sma-1* (encoding the cytoskeletal protein βH-spectrin) required for long tubule integrity [5, 6], and also regulates mRNA splicing [12] and synaptic transmission [13].

The Musashi family was also initially identified in *Drosophila*, where it is required for sensory bristle and neural development [14]. In mammals, MSI1 is predominantly expressed in neural precursor cells and multipotent CNS stem cells [15], while a second homologue, MSI2, is involved in the proliferation and maintenance of CNS stem cell populations [16]. MSI1 is up-regulated in a variety of human cancers, and negatively regulates Notch and Wnt pathway signaling via binding to Numb and APC, respectively [17, 18]. Knockdown of MSI1 function by means of siRNA [19], miRNA [20], or small-molecule inhibitors results in inhibition of tumor progression [21, 22]. In the nematode *C. elegans*, the sole family member MSI-1 is required for a specific step of male mating behavior and also acts via the Arp2/3 complex to mediate loss of memory [23, 24].

The ability of male *C. elegans* to mate efficiently requires tail flexures mediated by signaling by the neurotransmitter serotonin [25, 26], a step defective in *msi-1* mutants. Serotonin is a neurotransmitter involved in many *C. elegans* behaviors, including locomotion [27, 28], defecation [29], pharyngeal pumping [30, 31–33], egg-laying [29, 34, 35], and male tail flexure during mating [25, 26].

Since both serotonin and MSI-1 are required for male tail flexure, we have examined possible interactions between these two components. Findings here indicate that the human MSI1 inhibitor (-)-gossypol causes *C. elegans* male tail-turning defects, and this phenotype appears to be mediated by MSI-1. MSI-1 binds to and stabilizes mRNA encoding the G-protein alpha subunit GOA-1, which in turn alters expression of *tph-1* through *crh-1*/CREB. TPH-1 encodes the enzyme tryptophan hydroxylase that carries out the rate-limiting step of serotonin synthesis. In addition, EXC-7/HuR regulates serotonin level. By use of anti-serotonin staining, we found that mutants in either *exc-7, msi-1, goa-1, crh-1*, or *tph-1* show strong serotonin level decreases in neurons responsible for male tail flexure. These results demonstrate that a key EXC-7/MSI-1/GOA-1/serotonin pathway is responsible for this mating behavior in *C. elegans*. We believe that *C. elegans* could provide a useful model for understanding the interactions of these essential RNA-binding proteins in the regulation of neuronal signal transduction.

## Results

### Gossypol inhibits *C. elegans* MSI-1 interactions with mRNA

Once attracted to a hermaphrodite, the *C. elegans* male places its genital gubernaculum onto the hermaphrodite, and maintains contact while moving backward until sensory receptors in the tail locate the vulva near the center of the animal [36–38]. If the male reaches the end of the hermaphrodite before locating the vulva, the male tightly flexes its tail, maintaining contact, and curls around the hermaphrodite to move backward along the other side of the animal. Several passes may be required before the vulva is located [36]. The turns must be precise for the male tail to remain in contact, as the hermaphrodite may be moving during this process, so strong stimulation and flexure of the male diagonal muscles of the tail is necessary. The male’s ability to maintain tail contact with the hermaphrodite is defined as a “successful turn” (Fig. 1A, Supp. Video 1). When the male’s tail completely falls off the hermaphrodite, the turn is defined as “missed” (Fig. 1B, Supp. Video 2). An intermediate “sloppy turn” occurs when the tail loses contact briefly, but the male then reestablishes placement on the hermaphrodite (Fig. 1C, Supp. Video 3) [25].

**Figure 1.**
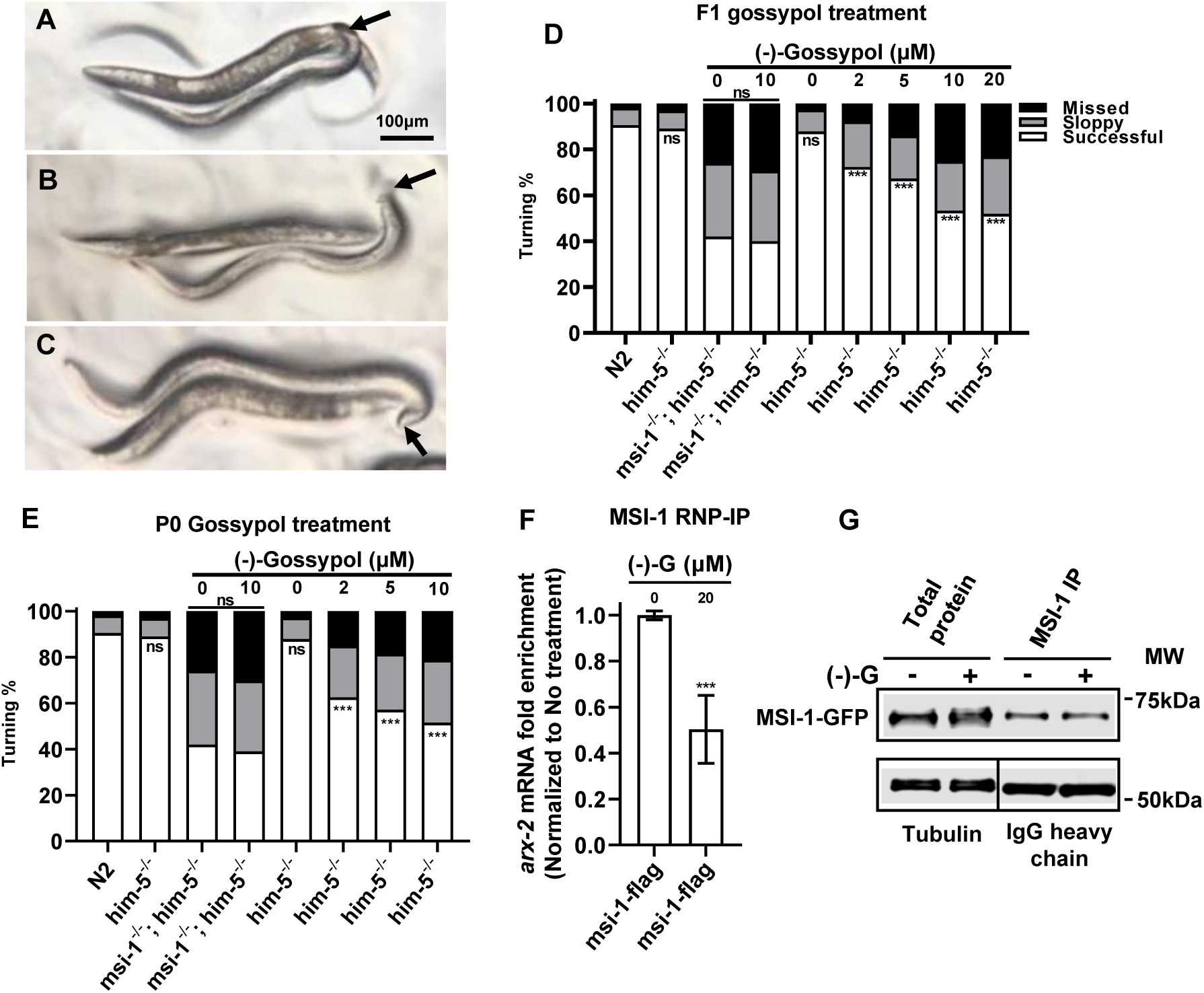
Human RNA-binding Musashi-1 inhibitor (-)-gossypol causes a *C. elegans* male mating defect. (A-C) Micrographs of male (smaller animal) backing up in attempting to find vulva on hermaphrodite (larger animal); arrows indicate gubernaculum on back end of male. Scale bar, 100 μm. **(A)** A “successful” turn, with male tail (arrow) maintaining contact while turning sharply to reverse direction of movement along the hermaphrodite. **(B)** A “missed” turn; the tail has fallen off the end of the hermaphrodite, and does not regain contact. **(C)** A “sloppy” turn; the male tail has slipped off the hermaphrodite, but through strong flexure, the male regains contact. **(D-E)** Gossypol feeding assays. The concentration of (-)-gossypol applied to each group is indicated at the top of the panel. Black bars represent percentage of missed turns, grey bars represent sloppy turns, and white bars represent successful turns. Numbers for N2 (wild-type) and *him-5^-/-^* control groups are re-used in multiple figures. NS, no significant difference; ***, P<0.001 (all from N2 except where bar is shown to indicate comparison). **(D)** F1 assay: Cultures of worms growing on a bacterial lawn were treated with the indicated concentrations of (-)-gossypol for 48-72 hr, and mating behavior of male progeny was measured. Total number of turns examined for each group ≥141. **(E)** P0 assay: L4 worms were treated with (-)-gossypol for 24 hr and the mating behavior of treated male animals was then tested. Total number of turns examined for each group ≥169. (Numbers for N2, *him-5^-/-^*, and untreated *msi-1^-/-^*;*him-5^-/-^* controls same as in Fig. 1D). (**F**)Quantitative PCR to compare the amount of *arx-2* mRNA bound by MSI-1 in the absence or presence of (-)-gossypol. (**G**)Western blot of RNP-IP of total protein and protein bound by MSI-1::Flag. Lysate and immunoprecipitated protein from both treated and untreated groups were loaded, and blotted for MSI-1::Flag and Tubulin.

The small molecule (-)-gossypol disrupts the interaction between mammalian Musashi1 and its target mRNAs [21]. To test whether this compound has the same effect on *C. elegans* MSI-1, we allowed *C. elegans* wild-type L4 hermaphrodites to feed and lay eggs in the presence of 0-20 μM (-)-gossypol, and scored the ability of F1 progeny males grown in this environment to turn during mating (Fig. 1D) (n.b.: We used a *him-5*(*e1450)* strain as wild-type; all strains in this study were crossed to this background to increase the frequency of males). At higher (-)-gossypol concentrations, the frequency of turning defects increased from less than 10% to approximately 50% (Fig. 1D), similar to the rate of mis-turning of *msi-1* mutants. Treatment of *msi-1*(*os-1*) mutants with (-)-gossypol showed no significant increase in the frequency of poor turns, consistent with (-)-gossypol action on *C. elegans* MSI-1 being a major mechanism of action of the drug for this behavior. These results indicate that (-)-gossypol induces the male turning defect and that (-)-gossypol likely functions by way of MSI-1. The effective concentration of (-)-gossypol on male turning (>= 10 μM for strongest effect) is also similar to that needed to affect human cell growth rates [21], which suggests that the drug gets into the nematode directly, rather than crossing the relatively impermeable cuticle [39].

As the effect described above could have resulted from defects in egg formation or early embryogenesis as well as direct effects on male behavior, we reduced exposure time of animals to the drug by placing L4 males on plates containing varying amounts of (-)-gossypol, and measuring mating behavior after just 24 hr (Fig. 1E). Again, application of (-)-gossypol substantially increased frequency of poor turns in wild-type males in a dosage-dependent manner, but had no effect on *msi-1* mutants. The rapid effect on turning is suggestive of action on a signaling pathway.

We next examined MSI-1 function. CRISPR/Cas-9 plasmids optimized for use in *C. elegans* were utilized to integrate *gfp* and 3 copies of *flag* DNA sequences at the 3’ end of the *msi-1* coding region [40]. Mating ability of male animals of this strain (BK561) has no significant difference compared to that of the parental *him-5* strain. (91.8% successful, 5.5% sloppy and 2.7% missed).

This *msi-1*::*gfp*::3x*flag* strain allowed determination of whether (-)-gossypol directly blocks MSI-1 binding to mRNA as it does in mammals. We performed ribonucleoprotein-immunoprecipitation (RNP-IP) with anti-Flag antibody in the presence or absence of (-)-gossypol, and measured via qRT-PCR the amount of target mRNAs bound (Fig. 1F). MSI-1 was previously reported to bind to the 3’UTR of *arx-2* mRNA [24]. Application of 20 μM (-)-gossypol interfered with MSI-1 function and lowered the amount of MSI-1-bound *arx-2* mRNA by 50% as compared to the amount bound in untreated controls. As control, a western blot (Fig. 1G) confirmed that MSI-1 levels were unchanged by (-)-gossypol treatment. We conclude that (-)-gossypol binds directly to MSI-1, and that this interaction disrupts the interaction between MSI-1 and downstream mRNA *arx-2*.

### MSI-1 binds *goa-1* mRNA to regulate male turning behavior

Male turning behavior is mediated by serotonin signaling [25, 26]. To study whether MSI-1 regulates this signaling, we performed an abbreviated screen of potential targets of MSI-1 binding to genes known to contribute to *C. elegans* male-turning behavior. Potential hits were identified via RNP-IP using anti-FLAG antibody to detect binding by MSI-1::GFP::3xFLAG. As negative control, GFP::3xFLAG was inserted (via CRISPR/Cas9 technology) into a protein that does not bind RNA, EXC-9 [41] (Fig. 2A). As *Flag* inserted at the endogenous *msi-1* locus was used instead of a construct overexpressing MSI-1::FLAG, we measured binding to *arx-2* mRNA as positive binding control for pulldown by MSI-1::FLAG. We found that *cat-1* mRNA and *goa-1* mRNA were pulled down even more strongly than was *arx-2* mRNA by MSI-1::FLAG (Fig. 2B).

**Figure 2.**
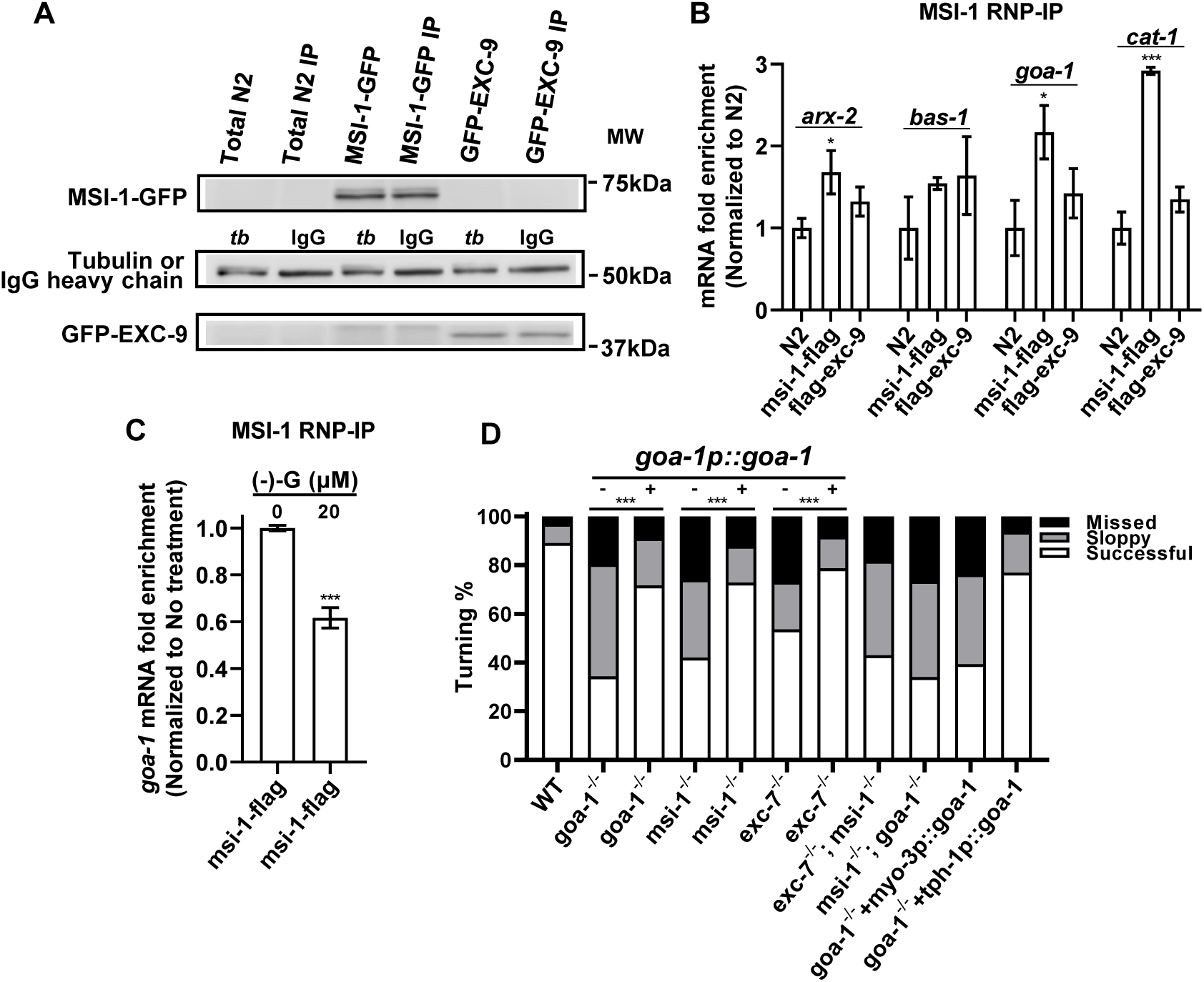
MSI-1 binds and stabilizes *goa-1* mRNA to regulate male turning behavior. **(A)** Western blot of MSI-1::Flag RNP-IP. Lysate and immunoprecipitated protein were loaded, and blotted for MSI-1::Flag, Tubulin, or IgG controls, and for Flag::EXC-9. **(B)** Comparison of amounts of specific mRNAs pulled down by MSI-1::3xFLAG from wild-type (N2), MSI-1-Flag, and Flag-EXC-9 cultures. *arx-2* is used as positive control. Statistical analysis was done via One-way Anova test. *, P<0.05; ***, P<0.001. **(C)** Quantitative PCR to compare the amount of *goa-1* mRNA bound by MSI-1 in absence or presence of (-)-gossypol. Statistical analysis was done via One-way Anova test. ***, P<0.001. **(D)** Male turning test of mutant strains, and the same strains containing a stable extrachromosomal array of *goa-1p::goa-1*. Bars as in Fig. 1: black, missed turns; grey, sloppy turns; white, good turns. ***, P<0.001. N (number of total turns)≥184. (Numbers for WT and untreated *msi-1^-/-^* controls same as in Fig. 1D).

The *cat-1* gene encodes an orthologue of human SLC18A1 and SLC198A2 (solute carrier family 18 members A1 and A2) and is predicted to have monoamine transmembrane transporter activity. The *goa-1* gene encodes an orthologue of the mammalian heterotrimeric G-protein alpha subunit Go (GNAO1). A previous transcriptome-wide RNA-binding analysis (CLIP-seq) for both endogenous and induced Msi1 in the intestinal epithelium identified mRNAs of GNAO1 and both SLC18A1 and SLC18A2 as likely targets of MSI1 [42]. In nematodes, *goa-1* mutants exhibited more severe male turning defects than those of *cat-1* mutants [25, 26]. We performed RNP-IP of MSI-1 in the presence or absence of (-)-gossypol, and utilized qPCR to measure binding to *goa-1* mRNA (Fig. 2C). In the treated group, the amount of *goa-1* mRNA bound was about 60% of that in the untreated control group, which confirmed that (-)-gossypol binding of MSI-1 affects the ability of this protein to bind to *goa-1* mRNA.

To see whether GOA-1 interaction with MSI-1 regulates serotonergic male turning behavior, we created a line overexpressing a stable multi-copy array of *goa-1* genomic DNA driven by its native promoter (Fig. 2D). Overexpression of *goa-1* caused some embryonic lethality (data not shown), but in surviving animals, *goa-1* expression substantially restored the turning ability of *goa-1(sy192)* mutant males, and in addition rescued the male turning defects of *msi-1* mutants (Fig. 2D). A *msi-1*; *goa-1* double homozygote shows no significant difference in rate of turning defects from those of *goa-1* homozygotes alone (P=0.2), consistent with these genes acting in a common pathway.

The *goa-1* gene is expressed in many tissues, including the male-specific diagonal muscles and neurons. As noted above, *msi-1* is expressed in many neurons. In order to determine whether GOA-1 is needed within the muscles or neural tissues to mediate male turning, we stably overexpressed genomic *goa-1* driven by either the *tph-1* promoter (expressing in the serotonergic neurons required for male turning) or the *myo-3* promoter (expressing in body wall muscles, including the diagonal muscles). Neuronal *goa-1* rescued male turning behavior in *goa-1* mutants, while *goa-1* expressed in muscles failed to rescue (Fig. 2D). These results indicate that *goa-1* functions in neurons to promote male turning ability.

### EXC-7 functions with MSI-1 to regulate male turning behavior

In human tissues, HuR binds to and stabilizes the 3’UTR of *Musashi1* mRNA [11]. Consistent with these observations, we found that in *C. elegans*, mutants in the *exc-7* gene (encoding the nematode homologue to human HuR) also exhibited male turning defects (Fig. 2D) (46% missed and sloppy turns), though at a lower frequency than did *msi-1* mutants (58% missed and sloppy turns, P=0.008 for the difference). We crossed *exc-7* to *msi-1* mutants to generate *msi-1; exc-7* double homozygotes. These animals show turning defects (57% missed and sloppy turns) no higher than those of *msi-1* homozygotes (P=0.1), with no evident synthetic effects. Stable *goa-1p::goa-1* overexpression in *exc-7* mutants partially rescues the *exc-7* phenotype (21% missed and sloppy turns) (Fig. 2D). These results indicate that EXC-7 likely acts upstream of *goa-1* to regulate its synthesis or activity.

### EXC-7, MSI-1, and GOA-1 regulate serotonin synthesis

Since serotonin is a key regulator of male turning behavior [25, 26], we tested whether *exc-7*, *msi-1*, and *goa-1* mutants were affected by exogenous serotonin. Mutants in these genes were soaked in buffer containing 1mM serotonin for 1 hour, and tested for male turning ability. For *exc-7*, *msi-1*, and *goa-1*, male turning defects were significantly rescued by exogenous serotonin (Fig. 3A), consistent with the three proteins functioning together in a pathway to regulate serotonin synthesis or signaling. Previous research discovered that GOA-1 affects transcription of the *tph-1* gene, which encodes tryptophan hydroxylase, the rate-limiting enzyme for serotonin biosynthesis, in both the HSN and NSM neurons in *C. elegans* [43]. Interestingly, GOA-1 activity inhibits *tph-1* transcription in HSN neurons, but promotes transcription in NSM neurons [43]. We examined TPH-1 function in male turning. Mutants in *tph-1* showed strong male turning defects (68% missed plus sloppy turns, Fig. 3A), and 1mM serotonin rescued these defects (18% missed plus sloppy turns). We tested the ability of *tph-1* genomic DNA (*tph-1p::tph-1*) to restore wild-type behavior to *exc-7, msi-1,* and *goa-1* mutants (Fig. 3B). Stable overexpression of *tph-1* genomic DNA rescued male turning ability to a similar level in each mutant (sum of missed plus sloppy turns: *exc-7*, 23%; *msi-1*, 30%; *goa-1*, 29%). We further confirmed a biological interaction between GOA-1 and TPH-1 by crossing the *goa-1* mutants to the *tph-1* mutants. The doubly homozygous mutant strain exhibited no significant difference in turning ability compared to either single mutant alone (Fig. 3C). These results suggest that TPH-1 acts downstream of EXC-7, MSI-1 and GOA-1, and that the activity of these proteins upregulates *tph-1* transcription.

**Figure 3.**
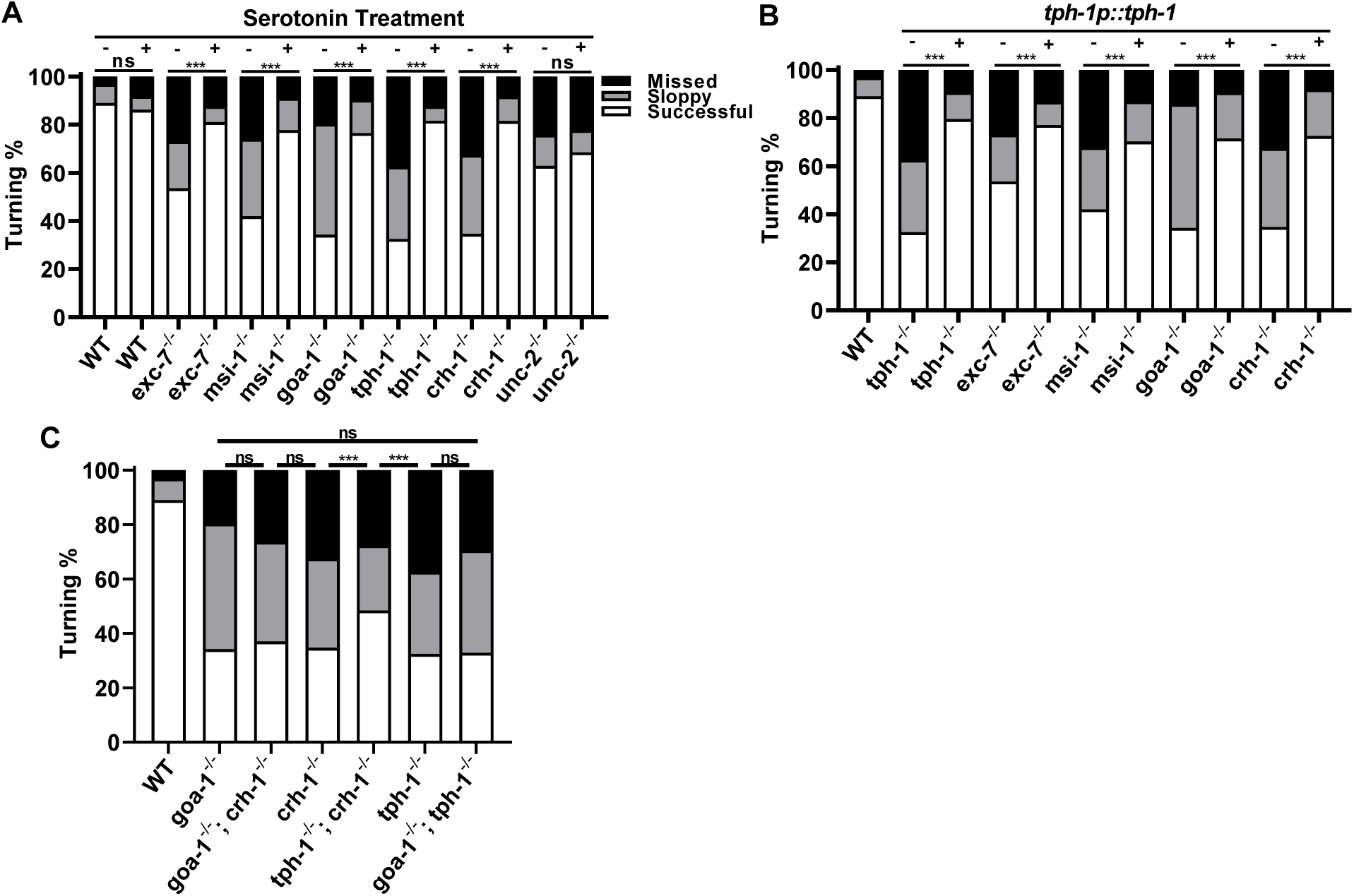
EXC-7 and MSI-1 regulate serotonin synthesis. **(A)** 1mM exogenous serotonin is applied (+) to various mutants. Black bars represent missed turns; grey bars sloppy turns, and white bars good turns (in all panels). N(number of total turns)≥169. **(B)** Male turning test with extrachromosomal stable array of *tph-1p::tph-1*. N(number of total turns)≥196. **(C)** Male turning test of doubly homozygous mutants of *goa-1*, *crh-1*, *tph-1*, and *unc-2*. with others. Statistical analysis is calculated via χ^2^ test. ***, P<0.001. N(number of total turns)≥124. In all panels, numbers for untreated WT and mutant controls same as in Fig. 2D.

Two major classes of protein are often involved downstream of G proteins in signaling: calcium channels and transcription factors [44]. Mutants in *unc-2* (encoding a calcium channel alpha subunit) and in *crh-1* (encoding a homologue of the cyclic AMP-response element-binding protein CREB) both affect serotonergic TPH-1 expression in other *C. elegans* neurons [45, 46]. Here, mutants in either gene showed male mating defects (Fig. 3A). The defects in *crh-1* mutants (65% missed plus sloppy turns) were comparable to those in the *tph-1* mutant, while *unc-2* mutants were significantly less impaired (37% missed plus sloppy turns). Defective turning in *crh-1* mutants can be rescued by soaking the animals in 1mM serotonin (Fig. 3A) or by overexpression of *tph-1p::tph-1* (Fig. 3B). Mutants in *unc-2*, however, could not be rescued by exogenous serotonin (Fig. 3A). This indicated that *unc-2* mutants, although exhibiting male turning defects, do not appear to act in the *goa-1*/*tph-1*/serotonin signaling pathway. We then crossed *crh-1*, *goa-1*, and *tph-1* mutants to each other (Fig. 3C). Double mutants that included the *goa-1* mutation were as strongly affected as the single *goa-1* mutant strain alone, consistent with GOA-1 acting together with CRH-1 and TPH-1. Finally, the *tph-1; crh-1* double homozygotes exhibited more successful turns than either mutation alone. We conclude that GOA-1 signals to CRH-1, which signals to TPH-1 to synthesize serotonin, although there may be more complicated regulatory mechanisms involved in the interactions of TPH-1 and CRH-1.

Although these results suggest that G-protein signaling affects serotonin synthesis to exert direct effects on neural function, it was also possible that the genes are involved in development of the male tail. We crossed these mutants to a strain carrying the serotonergic neuron marker construct *zdIs13*(*tph-1p*::*gfp*) and examined the adult male tails [47] (Fig. 4). For each of the single homozygotes mentioned (*exc-7*, *msi-1*, *goa-1*, *crh-1*, *tph-1*), the tails appeared to develop normally during the L4 larval stage, and the serotonergic neurons in the 1^st^, 3^rd^, and 9^th^ sensory rays on each side of the male tail showed normal expression from the *tph-1* promoter [48], so that male tail formation in these mutants appears to be normal.

**Figure 4.**
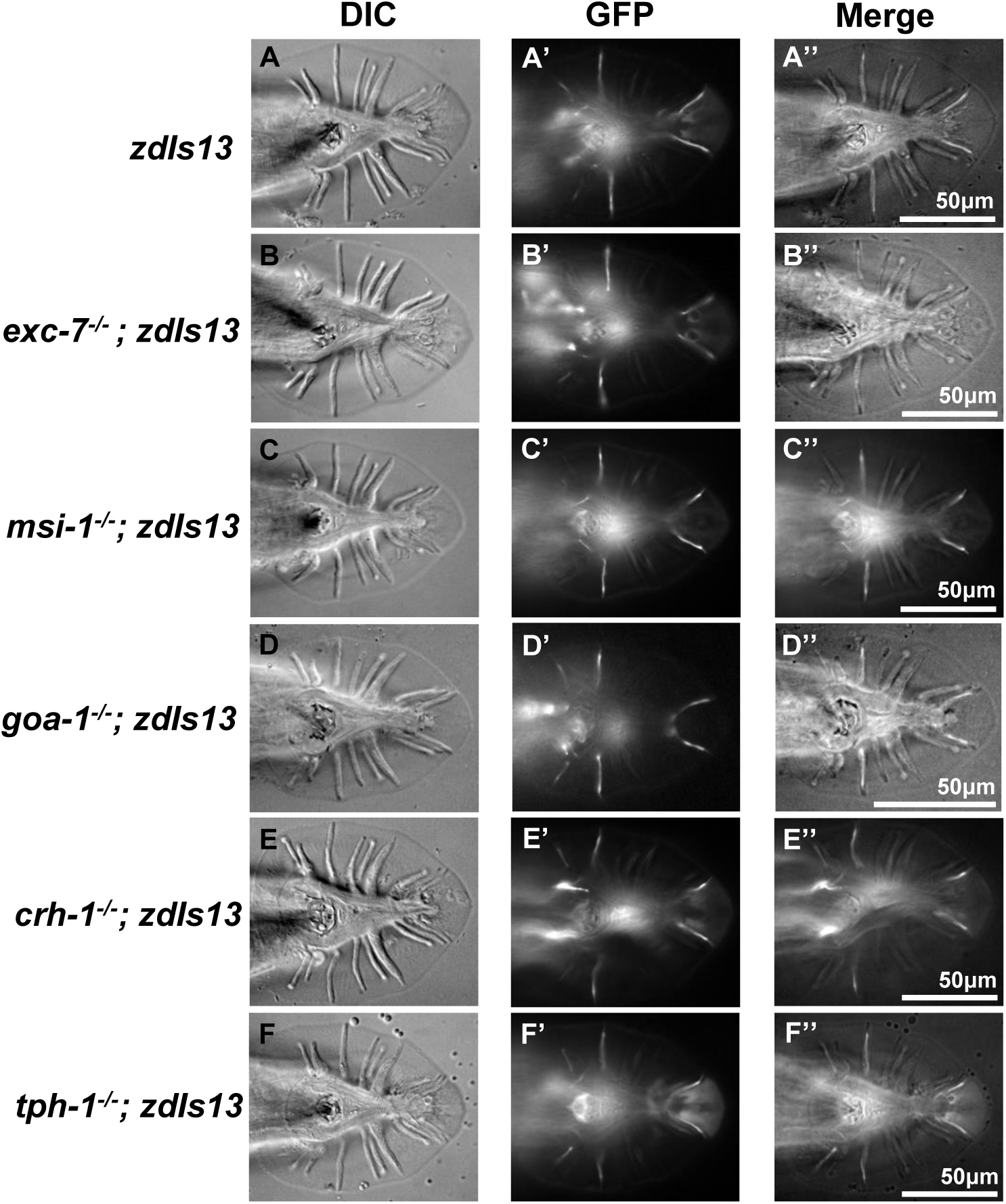
Male mutants’ tails show wild-type morphology. Micrographs of DIC, fluorescence, and DIC and fluorescence together of tail morphology of young adult males. Fluorescence marker is *zdIS13* (*tph-1p::gfp*). **(A)** wild type; **(B)** *exc-7^-/-^*; **(C)** *msi-1^-/-^*; (**D)** *goa-1^-/-^*; (**E)** *crh-1^-/-^*; (**F)** *tph-1^-/-^*.

### Serotonin levels vary in different mutants

There are several serotonin immuno-reactive neurons, including NSM (neurosecretory motor neuron), RIH (unpaired interneuron), ADF (amphid sensory neurons), AIM (interneuron), HSN (Hermaphrodite-Specific Neurons), RIH (unpaired interneuron), ADF (amphid sensory neurons), AIM (interneuron), RPAG (Right preanal ganglion neuron) in all animals, plus VC4 and VC5 motor neurons in hermaphrodites, and in males the bilaterally symmetric CP neurons 1-6 and sensory neurons inside rays 1, 3, and 9: R1B, R3B, and R9B, respectively [33]. The transcriptional reporter *zdIs13* expresses GFP under control of the *tph-1* promoter in these neurons [47], and was used to estimate relative levels of transcription of *tph-1* in different male-specific neurons and in the various mutant backgrounds tested in this study (Fig. 5B-H, complete results in Supp. Fig. 1A). Expression was highest in the CP5 and CP6 neurons, and decreased significantly in the more anterior CP neurons (Fig. 5B, 5H). Expression was also seen in ray neurons 3 and 9, and less strongly in ray neuron 1 (Fig. 5H). Expression in the mutant backgrounds was reduced in rough proportion to measured defects in male tail turning: Homozygous mutants in *exc-7* and in *msi-1* showed measurably less expression from the *tph-1* promoter, while *goa-1* mutants were even more strongly decreased (Fig. 5C-5G). These results were reinforced by experiments measuring serotonin levels directly through immuno-staining with anti-serotonin antibody (Fig. 5I-5Q, complete results in Supp. Fig. 1B). Serotonin levels were highest within the CP5 and CP6 neurons, and lowered in ray neurons (Note: As permeabilization of the animal to allow antibody entry required collagenase treatment, which degrades the male tail fan, the level of ray serotonin staining was measured only in ray neuron 1) (Fig. 5Q). In mutants, serotonin levels were decreased significantly in mutants affecting the RNA-binding proteins EXC-7 and MSI-1, while mutation in a *goa-1* or *crh-1* background were much more strongly affected (Fig. 5J-N, 5P). As expected, mutation of the serotonin synthase gene *tph-1* abolished serotonin staining (Fig. 5O, 5P). In the mutants, serotonin levels in mutants were more strongly affected than were levels of expression from the *tph-1* promoter; the magnitude of this effect may reflect use of an overexpression construct for measuring *tph-1* transcription. These results indicate that EXC-7, MSI-1, GOA-1, and CRH-1 promote serotonin synthesis, and that this regulation is partially achieved at the transcriptional level.

**Figure 5.**
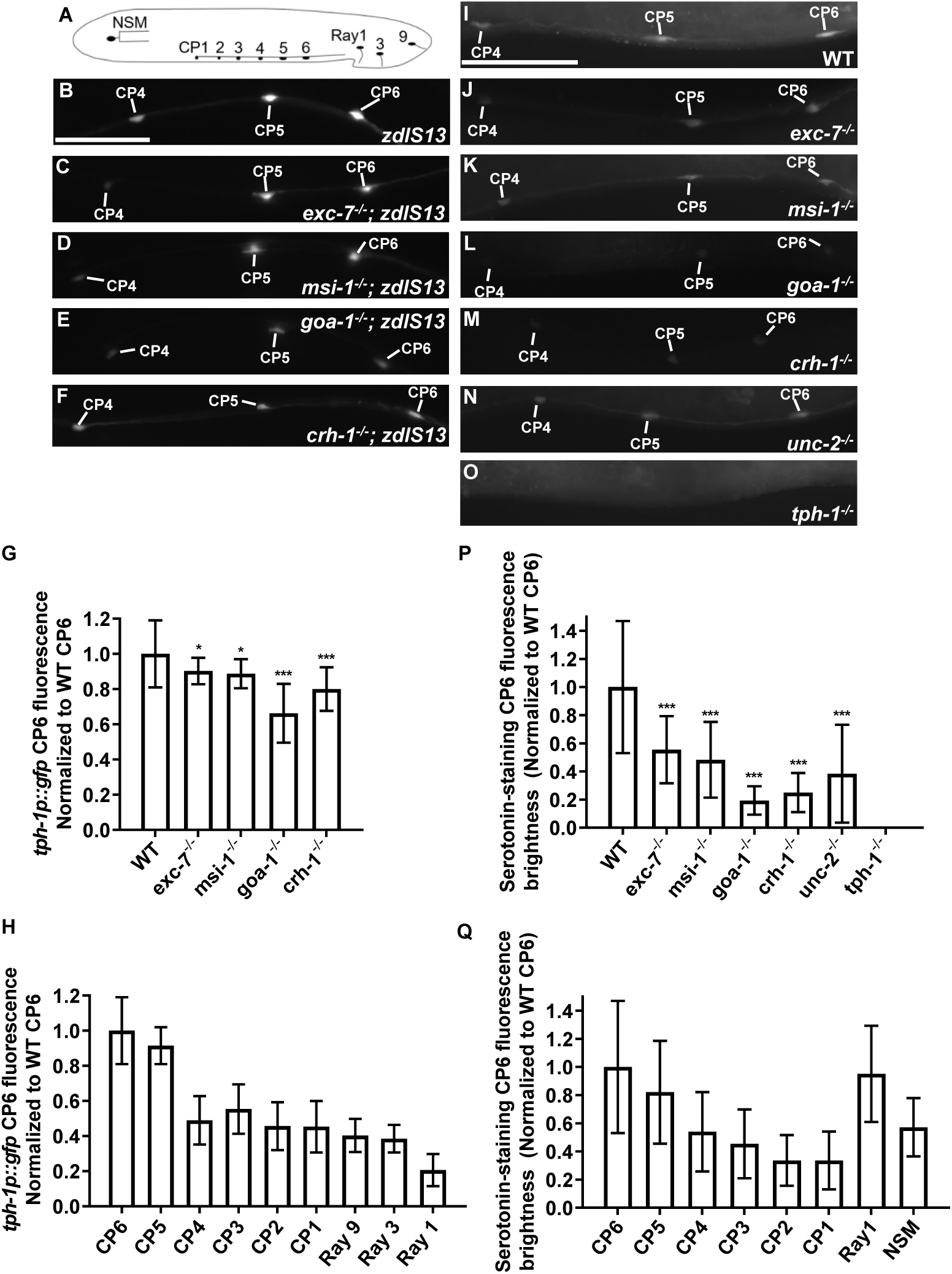
Anti-serotonin staining reveals different levels of endogenous serotonin in different mutants. **(A)** Diagram of serotonergic neurons compared in this study. **(B-F)** *tph-1* transcriptional reporter overexpression construct (*zdIs13*) expressed in strains of: **(B)** wild type; **(C)** *exc-7^-/-^*; **(D)** *msi-1^-/-^*; **(E)** *goa-1^-/-^*; **(F)** *crh-1^-/-^*. (**G**)Average expression of *tph-1* transcriptional reporter across male-specific neuronal CP6 cell bodies in wild-type and homozygous mutants of the indicated genes (all in *him-5* mutant background). (**H**) Relative fluorescence intensity of *tph-1* transcriptional reporter in neuronal cell bodies in wild-type animals. **(I-O)** Anti-serotonin staining of young adult males of genotype: **(I)** wild-type; **(J)** *exc-7^-/-^*; **(K)** *msi-1^-/-^*; **(L)** *goa-1^-/-^*; **(M)** *crh-1^-/-^*; **(N)** *unc-2^-/-^*; **(O)** *tph-1^-/-^*. (**P**) Relative average fluorescence intensity of neuronal cell CP6 bodies labeled via anti-serotonin staining for the tested strains, in *him-5* background. (**Q**) Relative fluorescence intensity in cells stained for serotonin in wild-type animals. *, P<0.05; ***, P<0.001.

### EXC-7, MSI-1, and GOA-1 are expressed in serotonergic neurons

In order to confirm that EXC-7, MSI-1, and GOA-1 regulate serotonin synthesis, we examined the cellular location of these proteins (Fig. 6). Previous research found that MSI-1 is expressed in the male-specific CP neurons [23]. We observed the *msi-1::gfp*::3xflag strain in which endogenous MSI-1 is fluorescently labeled, and confirmed that this protein is expressed in the cytoplasm of CP neurons and additionally in all nine of the male rays, as well as in the nerve ring, various head neurons, and male ray neurons (Fig. 6A, 6E). A strain carrying *mKate2* inserted into the endogenous locus of *exc-7* showed expression levels too low to detect reliably in male-specific neurons, but a line stably overexpressing *mCherry* driven by the *exc-7* promoter showed expression in the same CP neurons (CP1-CP6) as for *tph-1* expression, and in all nine male tail rays (Fig. 6B, 6B’, 6F). Finally, we sought GOA-1 location through insertion (via CRISPR/Cas9) of a fluorescent marker in the endogenous gene. Insertion of *mKate2* could not be used at the 5’ end of the coding region as it interfered with tethering of the protein to membrane, but the construct used, with *mKate2* insertion at the 3’ end of the coding region, rendered GOA-1 nonfunctional, as locomotion and animal shape resemble *goa-1* mutants. The expression pattern in the ventral cord and nerve rings, however, matches the location of this protein (Fig. 6C) [49]. In this strain, *goa-1* was not expressed in the CP neural cell bodies, as no obvious co-location pattern with *zdIs13* (*tph-1p::gfp*) was identified (Fig. 6D, D’, D’’). Expression of *goa-1* was strong in all male rays, however (Fig. 6G). We conclude that EXC-7, MSI-1, and GOA-1 mediate serotonin synthesis in the male tail rays, and that EXC-7 and MSI-1 potentially affect serotonin synthesis in the CP neurons.

**Figure 6.**
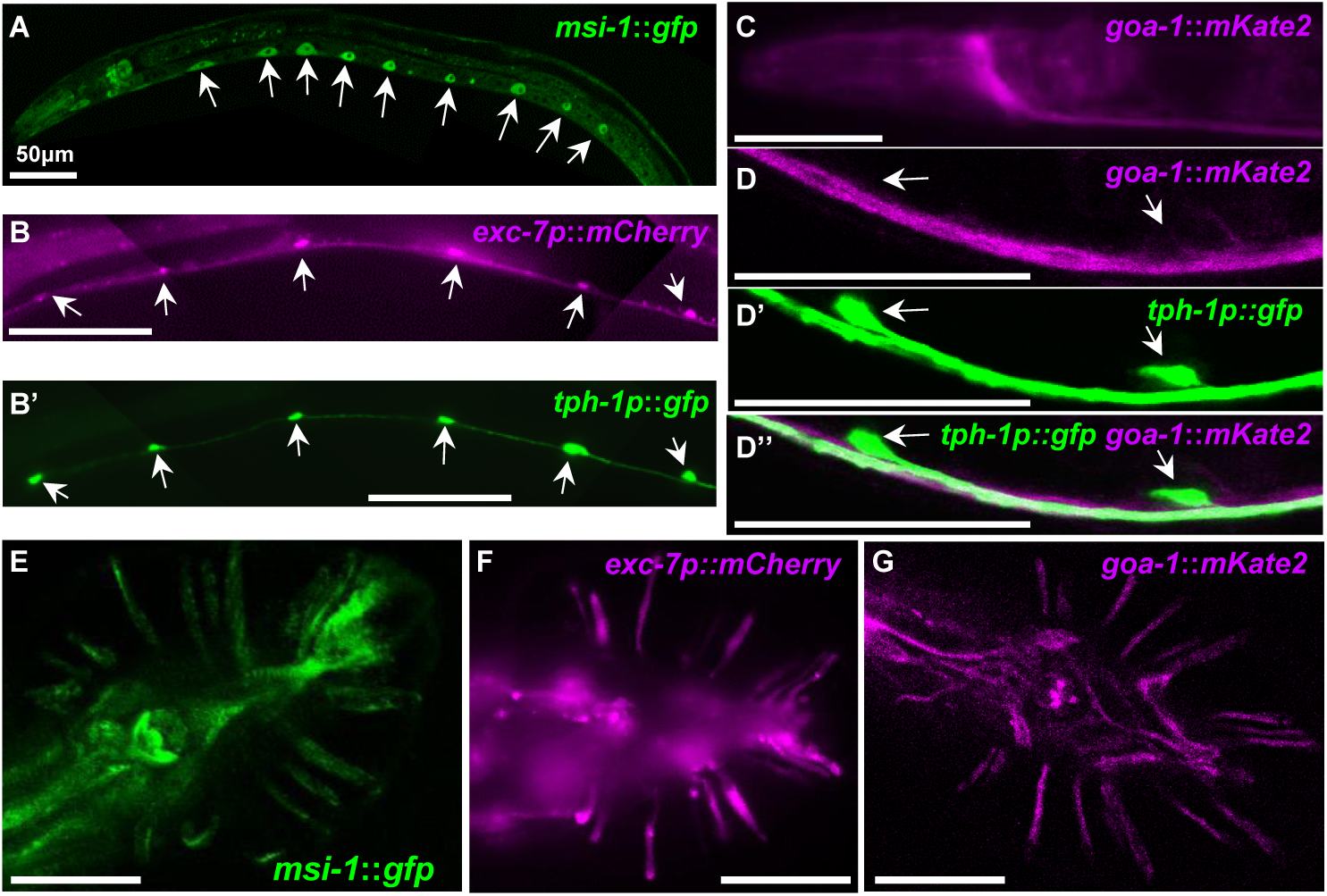
EXC-7, MSI-1, and GOA-1 are expressed in serotonergic neurons. **(A)** *msi-1*::*gfp* expression in cells of a young adult animal, including CP neurons (arrows). **(B-B’)** Co-expression in neuronal cell bodies and processes of CP neurons in a young adult male carrying: **(B)** *exc-7p::mcherry* overexpression; **(B’)** *tph-1p::gfp (zdIS13)* stable array. (**C**) *goa-1*::*mKate2* expression in nerve ring and ventral nerve cord. (**D**)*goa-1*::*mKate2* expression in CP neuron. **(D’)** *tph-1p::gfp (zdIS13)* expression in male CP5 and CP6 neuron. **(D’’)** Merge of *goa-1*::*mKate2* and *tph-1p::gfp (zdIS13)* at CP5 and CP6. (**E**) *msi-1*::*gfp* expression in male tail. (**F**) *exc-7p::gfp* expression in male tail. (**G**) *goa-1*::*mKate2* expression in male tail.

### Mutants exhibit CP neuron outgrowth defects

Serotonergic marker *zdIS13* expression of GFP became visible within the CP1-CP6 neurons at the L4 larval stage, with strong expression evident in the cell body and weaker expression in axons extending posteriorward along the ventral nerve cord (Fig. 7A). Previous ablation studies found that as few as two functional CP neurons were sufficient to provide wild-type male turning ability [25]. We examined axonal outgrowth of these mutants in strains carrying the *zdIs13* construct that strongly expressed GFP from the *tph-1* promoter (Fig. 7B-E). Defects observed included: a CP neuron either missing or not expressing GFP (Fig. 7B); some axonal outgrowth anteriorward instead of or in addition to posteriorward growth (Fig. 7C-D); and posterior axons not having fasciculated to the other CP axons (Fig. 7E). Surprisingly, a low level (2%) of otherwise wild-type animals carrying the *zdIs13* construct showed some CP axonal defects (Fig. 7F). The frequency of defects was exacerbated, to 5% in animals carrying homozygous defects in *exc-7*, and more strongly increased in animals with mutations in *msi-1*, *goa-1*, *crh-1*, or *tph-1* (7-17%). Growth of animals in plates containing 100μM (-)-gossypol also caused defects in the CP neurons (10%) (Fig. 7F). We conclude that genes responsible for serotonin synthesis affect CP neuron development, but other potential pathways are also involved that contribute to CP neural outgrowth.

**Figure 7.**
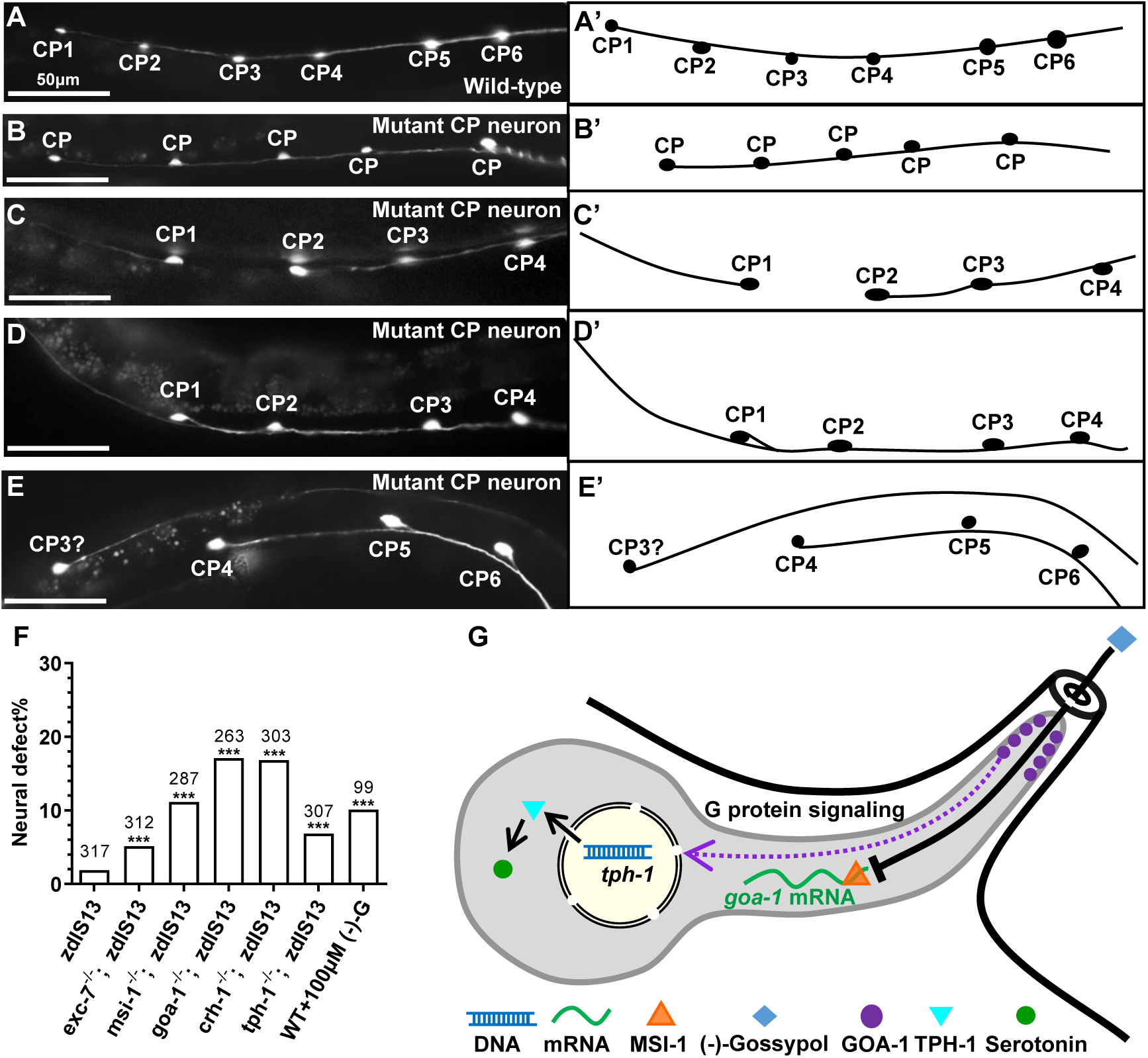
CP neurons exhibit various defects in mutants. **(A-E)**. Left-hand panels show fluorescence micrographs of CP neurons labeled via *zdIS13* [*tph-1p::gfp*]. Right-hand panels provide diagram of CP neurons 1-6 and axons as seen in left-hand panels. **(A, A’)** Wild-type neuronal pattern; axon of each neuron grows to the posterior (right). **(B, B’)** Animal with fewer labeled CP neurons (Missing or non-GFP-expressing CP). **(C, C’)** Animal with CP1 axon growing anteriorward instead of posteriorward. **(D, D’)** At least one CP neuron appears to send an axon both anteriorward and posteriorward. **(E, E’)** The process from the presumed CP3 neuron did not join cord of other CP neurons. (**F**) Graph showing frequency of animals exhibiting abnormal CP neuron morphology. (**G**) Model of MSI-1 influence on signal transduction machinery, and inhibition of synthesis by (-)-gossypol.

## Discussion

The results of this study delineate a novel and likely conserved pathway by which RNA-binding proteins mediate synthesis and signaling of the neurotransmitter serotonin. We found that (-)-gossypol, inhibitor of human Musashi, also binds to *C. elegans* MSI-1 to inhibit its binding to mRNA. In *C. elegans*, MSI-1 bound to *goa-1* mRNA to mediate male turning behavior; this binding could be interrupted by (-)-gossypol and cause defects in male turning. Mutations in *exc-7*, encoding the homologue of human HuR RNA-binding protein, also exhibited defects in male turning, as did homozygous mutations in the *goa-1* gene, encoding the alpha subunit of the Go heterotrimeric G-protein signal transduction complex. Stable expression of *goa-1*, in contrast, rescued the turning defects in both homozygous *msi-1* and homozygous *exc-7* mutants.

Two other genes, *crh-1* and *tph-1*, acted downstream of *goa-1*; both encode proteins that regulate production of serotonin [43, 45, 46]. CRH-1 is the homologue of cyclic AMP-response element-binding protein CREB, while TPH-1 tryptophan hydroxylase enzyme conducts the rate-limiting step in serotonin synthesis [50]. Mutations in these genes also caused male turning defects. All of these mutants showed reduced levels of serotonin in male ray and CP neurons, and application of serotonin to affected animals rescued mutants in all of the genes studied.

Mating behavior of nematode males is stimulated by the neurons in the rays of the male tail, which have small openings to allow the ray neurons to detect the hermaphrodite by means of chemotactic and thigmotactic signaling [36, 37]. As treating males for a relatively short time in (-)-gossypol at concentrations in the same range as for human cells in culture strongly affected turning ability, we infer that the male ray openings are the most likely entry site for (-)-gossypol to affect male turning, and that the ray neurons are the most likely site of action of the pathway described above.

A model to summarize serotonin regulation at male tail rays is presented in Fig. 7G. In wild-type animals, MSI-1 binds to *goa-1* mRNA to promote its translation. GOA-1 upregulates *tph-1* transcription to facilitate serotonin synthesis through CRH-1. EXC-7 is also involved in this pathway to upregulate *tph-1* expression. The inhibitor (-)-gossypol passes through the ray opening to gain access to the neuron and interfere with MSI-1 function, leading to decreases in the amount of *goa-1* mRNA and concomitant decrease in serotonin synthesis.

Male ray neurons synapse with the male-specific CP neurons [51]. The CP neurons are not well-studied, but appear to innervate the diagonal muscles needed for male tail flexure. Interestingly, while all of the genes studies were found to be expressed in Rays 1, 3, and 9, the GOA-1 protein did not appear to be expressed in the CP neurons (although we could not exclude the possibility that GOA-1 is found solely at synapses to ray neurons). This reinforces our conclusion that the ray neurons are the site of (-)-gossypol action on serotonin synthesis.

All of the proteins in the pathway have been well conserved from nematodes to humans, and several are involved in human diseases. In particular, the human homologues of nematode EXC-7 (HuR), MSI-1 (MSI1 and MSI2) and GOA-1 (GNAO1) have been implicated both in disorders that affect serotonin signaling, and in human cancers. Upregulation of MSI and Hu proteins is seen in multiple tumor tissues. siRNA-mediated knockdown of these genes slows tumor cell growth, as does treatment with (-)-gossypol or other inhibitors of these RNA-binding proteins [10, 19–22].

Mutations in GNAO1 have been linked to various human diseases, including breast carcinoma [52], lung adenocarcinoma [53], as well as epileptic encephalopathy [54, 55], and movement disorder [56–58]. Levels of GNAO1 are also significantly decreased in individuals with schizophrenia [59]. Finally, GNAO1 is predominantly expressed in brain, pancreas, and the gastrointestinal tract, all tissues that produce serotonin.

A pathway regulating serotonin synthesis from RNA-binding proteins to G-protein signaling to serotonin synthesis appears to be conserved from nematodes to humans. (-)-Gossypol, which inhibits the action of these RNA-binding proteins in humans exerts a similar effect on a serotonergic pathway in humans. Unlike many drug effects on *C. elegans*, the impermeability of the cuticle does not appear to be a hindrance to the action of (-)-gossypol on male tail turning [60, 61]. Assays to gauge male turning ability in *C. elegans* therefore presents a non-invasive, useful method for gauging the effectiveness of (-)-gossypol and other drugs on HuR and MSI function both to regulate serotonin levels, and possibly to affect growth rates of tumor cells.

## Supplementary Files

1. “Supplementary Figure 1” – Graphs of complete serotonin levels within individual neuronal types.

2. “Supplementary Data” includes: Supplementary Table 1 (Nematode strains used), Figure Legend for supplementary Figure 1, and description of Supplementary Videos

### Supplementary Figure Legend 1: Measured serotonin levels

(A) Immunostaining with anti-serotonin antibody. Brightness levels were normalized to the brightness of the CP6 neuron.

(A) *tph-1p*::*gfp* reporter gene, normalized to brightness of the CP6 neuron.

### Supplementary Video Legends

3. “SuppVideo 1-Successful Turn”: Male *C. elegans* adult (∼ 1mm in length) backs along an adult hermaphrodite, successfully maintaining close contact of its gubernaculum (at the posterior end of the male) with the hermaphrodite at all times, including while reversing direction, in order to locate the hermaphrodite’s vulva.

4. “SuppVideo 2-Missed Turn”: Male *C. elegans* adult backs along an adult hermaphrodite, but fails to turn at the end of the hermaphrodite and continues backing past the hermaphrodite tail.

5. SuppVideo 3-Sloppy Turn”: Male *C. elegans* adult backs along an adult hermaphrodite, fails to turn successfully at the appropriate time, but is able to turn its tail enough to regain contact of its gubernaculum with the hermaphrodite in order to try again.

## Acknowledgements

We thank David H. Hall (Albert Einstein College of Medicine) for information on CP neurons. Some strains were provided by the Caenorhabditis Genetics Center (CGC), which is funded by NIH Office of Research Infrastructure Programs (P40 OD010440). ZY was supported by Kansas University Graduate Research Fund #2144091. This study was supported in part by National Institutes of Health grants R01 CA178831, CA191785, and CA243445 (to LX), and Kansas Bioscience Authority Rising Star Award (to LX).

## Methods

### C. elegans strains

*C. elegans* strains were grown on BK16 bacteria (a streptomycin-resistant derivative of OP50) at 20C according to standard techniques [62]. All mutants were crossed to *him-5*(*e1490*) to create strains with a high proportion of males [63]; this mutation is present in all strains used here (including wild-type) though is not listed in tables or figures. All strains used are listed in Supplementary Table 1.

### DNA constructs and plasmids

*goa-1* driven by its native promoter was amplified from N2 wild-type genomic DNA using forward primer 5’-GTATTCAAAAGTTTCGCGCCAATGCG-3’ (3136 bps upstream of ATG) and reverse primer 5’-GTACGGTAGTCCCATATGCAATTTCCC-3’. *tph-1* driven by its native promoter was amplified from N_2_ wild type genomic DNA using forward primer 5’-GCAATACTATTTTTCGGTGGTCTTCCC-3’ (3139 bp upstream of ATG) and reverse primer 5’-CGTGTCACATCCTTTATGTGCGTTG-3’. pBK263 (*tph-1p::goa-1*) was derived from pCVO1. For this construct, the *tph-1* promoter was amplified with forward primer 5’-CTTGGAAATGAAATAAGCTTGCATGCGCAATACTATTTTTCGGTGGTCTTCC-3’ (3139 bp upstream of ATG) and reverse primer 5’-CTTCCTGTGACATGGTACAACCCATATGATTGAAGAGAGCAATGCTAC-3’. Gibson assembly was carried out to assemble pBK263 from NEB NEBuilder High-Fidelity Master Mix (Catalog #: M5520). pBK272 (*myo-3p::goa-1*) was derived from pCVO1. The *myo-3* promoter was amplified with forward primer 5’-CTTGGAAATGAAATAAGCTTGCATGCCTGTTTGATGAAAACCAATGAAACAAGTG- 3’ (2525 bp upstream of ATG) and reverse primer 5’-CTTCCTGTGACATGGTACAACCATTTCTAGATGGATCTAGTGGTCGTG-3’. For both of these last constructs, the *goa-1* coding region insertion was amplified with forward primer 5’-GTAACCGGTATGGGTTGTACCATGTCACAGG-3’ and reverse primer 5’-CTACGGCCGCAAATAAGATATCAATGAGTGGGTGCAGTC-3’ and restriction enzyme AgeI (NEB Catalog #: R3552S) and EagI (NEB Catalog #: R3505S).

### Male turning assay

We adopted the male turning assay developed in the Loer laboratory [25]: L4 males were picked and adapted to fresh 60mm plates for 24 hours, then scored via a double-blind assay. Individual adult males were placed on a 30mm plate with one drop of bacteria and 10-12 adult *unc-51* (paralyzed) young adult hermaphrodites. Each male was observed attempting to mate, and 5 turns (technical replicates) of the male backing along the hermaphrodite were scored (fewer if the animal had movement defects, e.g. for *unc-2* mutants). Each turn was categorized as either “good,” “sloppy,” or “missed”. At least 34 males (biological replicates) were tested for each genotype measured, with the exact numbers listed in Figshare. In some cases, the same numbers for controls are presented in multiple figures.

### Microscopy

Nematodes were examined via a Zeiss microscope equipped for both epifluorescence and DIC microscopy (Carl Zeiss, Thornwood, NY) and photographed by use of an Optronics MagnaFire Camera. Animals were placed on 3% agarose pads in M9 solution + 35mM NaN_3_. Images of CP neurons of larger worms (Fig. 7B, C) required four photographs that were “stitched” together to provide a picture of the six CP neurons. Contrast on DIC images was uniformly enhanced over the entire image to increase clarity.

Subcellular location of fluorescent proteins at high resolution was examined through a FluoView FV1000 laser-scanning confocal microscope (Olympus, Tokyo, Japan). Lasers were set to 488 nm excitation and 520 nm emission (GFP), or 543 nm excitation and 572 nm emission (mKate2). All images were captured via FluoView software (Olympus) and collocation was analyzed by use of ImageJ software.

For categorizing frequency of neural outgrowth defects, individual animals (number indicated on graph, biological replicates) were examined for presence of any abnormalities in CP neuron trajectory, and classified as “normal” or “abnormal.”

### Ribonucleoprotein-Immunoprecipitation, qPCR and western blot

The RNP-IP technique was adapted as previously described [24, 64]. Ribonucleoside Vanadyl Complex was from NEB (Catalog #: S1402S). Anti-FLAG® M2 Magnetic Beads was from Sigma-Aldrich (Catalog #: M8823). RNA was reverse-transcribed with random primers from Applied Biosystems (Catalog #: 4374966). qPCR was performed on 3 duplicates (technical replicates) for each of the samples tested, through use of the SYBR kit from Applied Biosystems (Cat #: A25742). Expression levels were normalized to the nematode housekeeping gene *tba-1* (encoding α-tubulin). Western blot was performed with Anti-Flag antibody DYKDDDDK Tag Monoclonal Antibody (FG4R) from Fisher Scientific (Catalog #: MA1-91878). Monoclonal Anti-α-Tubulin antibody produced in mouse was applied from Sigma-Aldrich (Catalog #: T5168).

### Anti-serotonin staining and measurement

Anti-serotonin staining protocol is adopted from that of Loer *et al.* [25]. Anti-serotonin antibody produced in rabbit (Catalog #S5545) and secondary antibody Anti-Rabbit IgG (whole molecule)–TRITC antibody produced in goat (Catalog #T6778) were from Sigma-Aldrich. ImageJ was used to measure the brightness of serotonin in tissues in the red channel; some animals carried the *zdIS13*(*tph-1p*::*gfp*) construct, which allowed measurement of TPH-1 transcriptional expression levels in the green channel.

### Statistics

For male tail-turning assays, statistical differences between groups of animals were calculated via χ^2^ test of good, missed, and sloppy turns against the control results for N2 or *him-5*^-/-^ animals. For serotonin staining, at least 30 animals (biological replicates) were photographed for each fluorescence measurement, and results averaged. Since collagenase treatment was needed to dissolve the cuticle, not all animals showed serotonin staining. Such animals were not used for calculating brightness of neural staining, except for *tph-1* mutants, which exhibited no staining in any animal. P-values of anti-serotonin staining and *tph-1::gfp* brightness are calculated via one-way ANOVA.

Complete data (including total number of turns examined) and calculated P-values results are included on Figshare.

## Data Availability

Complete data with statistics of male turning assays are at https://figshare.com/s/c00d7af4979004b1538c.

Serotonin staining data and measurements of fluorescence are at https://figshare.com/s/8bade09b55cbb74abf33.

Fluorescently labelled nematode strains will be submitted to the *Caenorhabditis* Genetics Center, or (if not accepted) will be available from the corresponding author.

## Author Contributions

ZY helped design, and performed all genetic and staining experiments, and helped write the paper. LL and ZY performed biochemical experiments. XQ helped in experimental design. LX contributed expertise on drug usage, and designed biochemical experiments. MB helped design genetics experiments, oversaw the project, and wrote the paper.

---

**Supplementary Figure 1.**
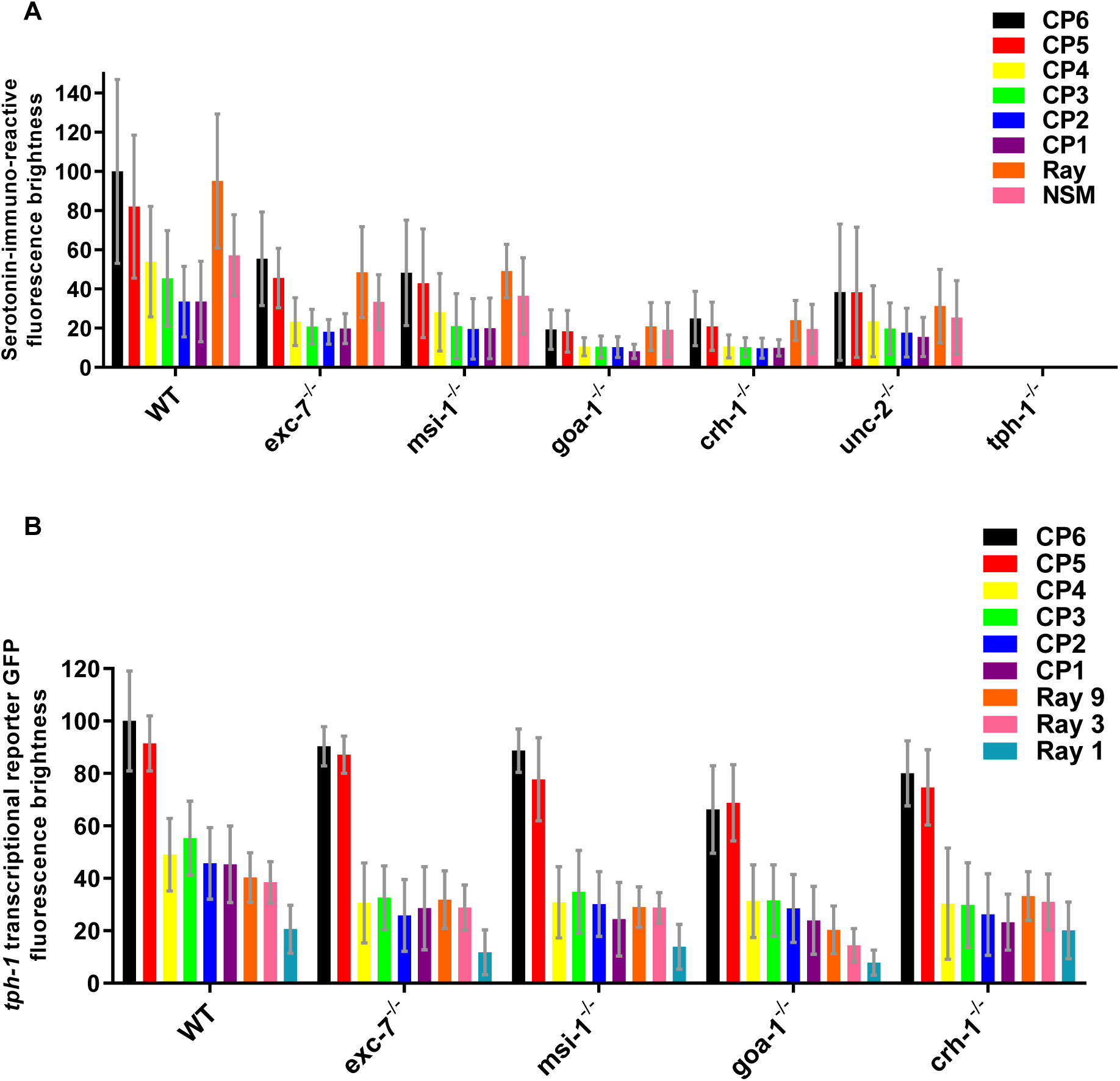
Measured serotonin levels. **(A)** Immunostaining with anti-serotonin antibody. Brightness levels were normalized to the brightness of the CP6 neuron. **(B)** *tph-1p::gfp* reporter gene, normalized to brightness of the CP6 neuron.

